# Induction of autophagy by trehalose limits opportunistic mycobacterial infections in HIV-infected macrophages

**DOI:** 10.1101/202697

**Authors:** Vartika Sharma, Muzamil Makhdoomi, Purnima Kumar, Nabab Khan, Sarman Singh, H N Verma, Kalpana Luthra, Sovan Sarkar, Dhiraj Kumar

## Abstract

Opportunistic bacterial infections amongst HIV-infected individuals pose serious health challenge. While immediate control of bacterial pathogens is typically attributed to innate defense mechanisms, whether HIV-mediated modulation of innate mechanisms like autophagy promote opportunistic infections, remains obscure. Using U1.1 and U937 macrophages, we show, HIV activation or infection inhibits autophagy and helps survival of pathogenic *Mycobacterium tuberculosis* and non-pathogenic non-tuberculous mycobacterial strains (NTMs) like *Mycobacterium avium* complex and *Mycobacterium fortuitum*. HIV achieves this by blocking xenophagy flux, which could be reversed by the autophagy inducer trehalose that kills intracellular *Mtb* and NTMs. We found trehalose acts as a PI (3,5) P_2_ agonist and activates TRPML1 to induce autophagy. Remarkably, trehalose treatment significantly reduced p24 levels in PBMCs infected with clinical HIV strains and in PBMCs derived from treatment-naive HIV patients. Taken together, our study highlights the immense potential of autophagy modulators in the therapeutic intervention of HIV and associated opportunistic infection.

## Introduction

Opportunistic infections in human immunodeficiency virus-1 (HIV-1) infected individuals - are one of the major causes leading to Acquired Immunodeficiency Syndrome (AIDS) associated mortality. The CD4^+^ T lymphocytes are major targets for HIV-1 infection. The decline in CD4^+^ T cells is directly correlated to the increased viral load and is used as a clinical benchmark to establish the onset of AIDS^1^. However, several opportunistic bacterial infections, which aggravate the clinical presentation of AIDS patients, do not require a strong T cell-response initially for their effective containment by the host-immune system. These bacterial infections rather require a strong innate defense response to get controlled, in absence of which, they could establish a successful infection. It is therefore critical to understand whether and how innate defense mechanisms are also perturbed by HIV-1 infection. That the cells of myeloid lineage like monocytes, macrophages and dendritic cells can get infected with HIV-1 and also serve as reservoirs, underscores the importance of innate defense mechanisms in the clinical presentation and management of AIDS^2, 3^. Several studies in the past suggest HIV-1 mediated perturbation of host innate immune system including those involving surfactants and effector cell functions like phagocytosis, efferocytosis and response to inflammatory stimuli like TNFs ^4^. However, there are also studies, which did not report any defect in alveolar macrophage function among HIV-1 infected individuals unless they were also tobacco smokers^5^.

Tuberculosis (TB), which infects a large section of population, especially in the endemic countries, remains as an asymptomatic infection in more than 90% of cases. However, among HIV-AIDS patients, nearly 40% of them may show active TB^6^. Similarly, certain non-tuberculous mycobacterial strains (NTMs) like *Mycobacterium avium, Mycobacterium fortuitum*, *Mycobacterium kansasi*, *Mycobacterium terrain* and *Mycobacterium intracellulare*, most of which are largely non-pathogenic in humans, may cause active disease among HIV-1 infected individuals ^7^. HIV-1-TB coinfection as co-morbidity has been studied extensively, and features like increased HIV-1 replication at the site of co-infection, HIV-1 induced killing of CD4^+^-T cells in the granulomas, prevent *Mtb* killing by manipulating macrophage function and functional changes in *Mtb*-specific T cells has been reported previously^8^.

Among various anti-bacterial innate defense mechanisms, autophagy has emerged as a central player in the recent times. Several pathogenic bacterial species including *Mycobacterium tuberculosis, Salmonella typhimurium, Legionella* sp., *Listeria* etc. are known to perturb the host autophagy machinery in order to survive within the host cells^9^. Interestingly, HIV-1 infection is also known to inhibit the final stage of autophagy i.e. fusion of autophagosomes with the lysosomes for maturation into auto-lysosomes. The rate of autophagic cargo degradation through autophagosome maturation is described by autophagy flux, which compares cargo accumulation in the presence of a lysosomal function inhibitor with respect to control. Importantly HIV-1 infection triggers early stages of autophagy leading to formation and accumulation of more autophagosomes^10–12^ and more virion biogenesis as well. Despite both HIV-1 and other bacterial infections perturbing the host autophagy pathway, very little is known about whether within the same host, autophagy perturbation by a viral infection could inadvertently help other bacterial infections. There are very limited systematic studies, which connect the high opportunistic bacterial infections with HIV-1 induced perturbation of innate defense mechanisms, especially autophagy, in the host. Few studies, which somewhat explored this exciting crosstalk reported a role of Vitamin D_3_ mediated induction of autophagy to control both HIV-1 replication and *Mycobacterium tuberculosis* (*Mtb*) survival within macrophages, either singly or during co-infections^13–15^. However there is no such study yet that explores whether HIV mediated impairment of autophagy could promote opportunistic infection of *Mtb* or other NTMs during HIV co-infection.

Modulation of autophagy can influence xenophagy, as has been shown for several small molecule autophagy modulators like rapamycin^16, 17^. A potent, safe autophagy inducer is the naturally occurring disaccharide trehalose. Trehalose induces autophagy independently of the mechanistic target of rapamycin (mTOR) pathway and was first reported to facilitate aggrephagy (clearance of aggregation-prone proteins) in mammalian and neuronal cells^18^. The exact mechanism of trehalose-mediated induction of autophagy however remains debatable, despite some recent developments like its role as competitive inhibitor of glucose transporters (GLUT) and the identification of GLUT8 as mammalian trehalose transporter^19–21^. Interestingly, virulent *Mtb* is known to secrete trehalose outside of the cell, which is a by-product of the enzymatic machinery that synthesizes complex glycolipids for mycobacterial cell wall ^22^.

In this study, we show that active HIV-1 replication either through reactivation or productive infection results in inhibition of autophagy flux and accumulation of autophagosomes. More importantly, the HIV-1 mediated block in autophagy flux, we show for the first time, helps enhance the survival of *Mtb* as well as non-tuberculous mycobacterial strains in HIV-1 infected macrophages. Using trehalose, we show that autophagy induction not only controlled *Mtb* survival alone or when co-infected with HIV-1, but could also control p24 levels in PBMC cultures. Moreover, we report here a previously unrecognized mechanism of action of trehalose for inducing autophagy in macrophages.

## Results

### Activation of HIV-1 replication in macrophages results in impaired autophagy flux

To understand the effect of HIV-1 replication on macrophage autophagy response, we used U937 and U1.1 monocytic cell systems. U1.1 cell line is a subclone of U937 cells, which is chronically infected with HIV-1 and contains an integrated copy of HIV-1 in its genome. This cell line shows a minimal constitutive viral expression. The viral expression can be induced upon treatment with certain cytokines and PMA^23^. In both U937 and U1.1 monocytes, bafilomycin A_1_ (BafA_1_) treatment resulted in the accumulation of LC3II, indicating basal autophagy flux in these cells (Fig 1A). Treatment with PMA, which is known to activate these monocytes into macrophages, had no considerable effect on autophagy flux in U937 cells (Fig 1B). However in U1.1 cells PMA treatment led to increased LC3II protein even in the absence of BafA_1_, which did not increase further upon BafA_1_ treatment, suggesting a block in autophagy flux (Fig 1B). A time-course immunoblot analysis of p24 in U1.1 cell lysates upon PMA treatment confirmed activation of HIV-1 replication by 12 hours, which reached to its peak levels by 48 hours and was expressed at high level even till 96 hours post-PMA treatment (Fig. 1C). Decrease in autophagy flux upon PMA dependent HIV-1 activation in U1.1 cells was also confirmed by flow cytometry, where cells were stained for detergent insoluble LC3II intensity in untreated, PMA-treated or PMA followed by chloroquine-treated cells (Fig 1D). In agreement with the immunoblot data, U937 cells showed autophagic flux in both untreated and PMA stimulated cells (Fig 1D). Finally we analyzed autophagosome maturation via the pH-sensitive tandem-fluorescent-tagged autophagy reporter, mRFP-EGFP-LC3, which can distinguish between the autophagosomes (mRFP^+^/EGFP^+^) and the autolysosomes(mRFP^+^/EGFP^-^) because EGFP is quenched in the acidic environment of the latter compartment^24,25^. In U1.1 monocytes, BafA_1_ prevented autophagosome maturation as evident by increased autophagosome/autolysosome ratio (ratio of mRFP^+^/EGFP^+^ to mRFP^+^/EGFP^-^ puncta), which remained unchanged in U1.1 cells treated with PMA in the absence or presence of BafA_1_ (Fig 1E). Taken together, these data suggest that activation of HIV-1 replication in macrophages results in the impairment of autophagic flux in macrophages.

**Figure 1:**
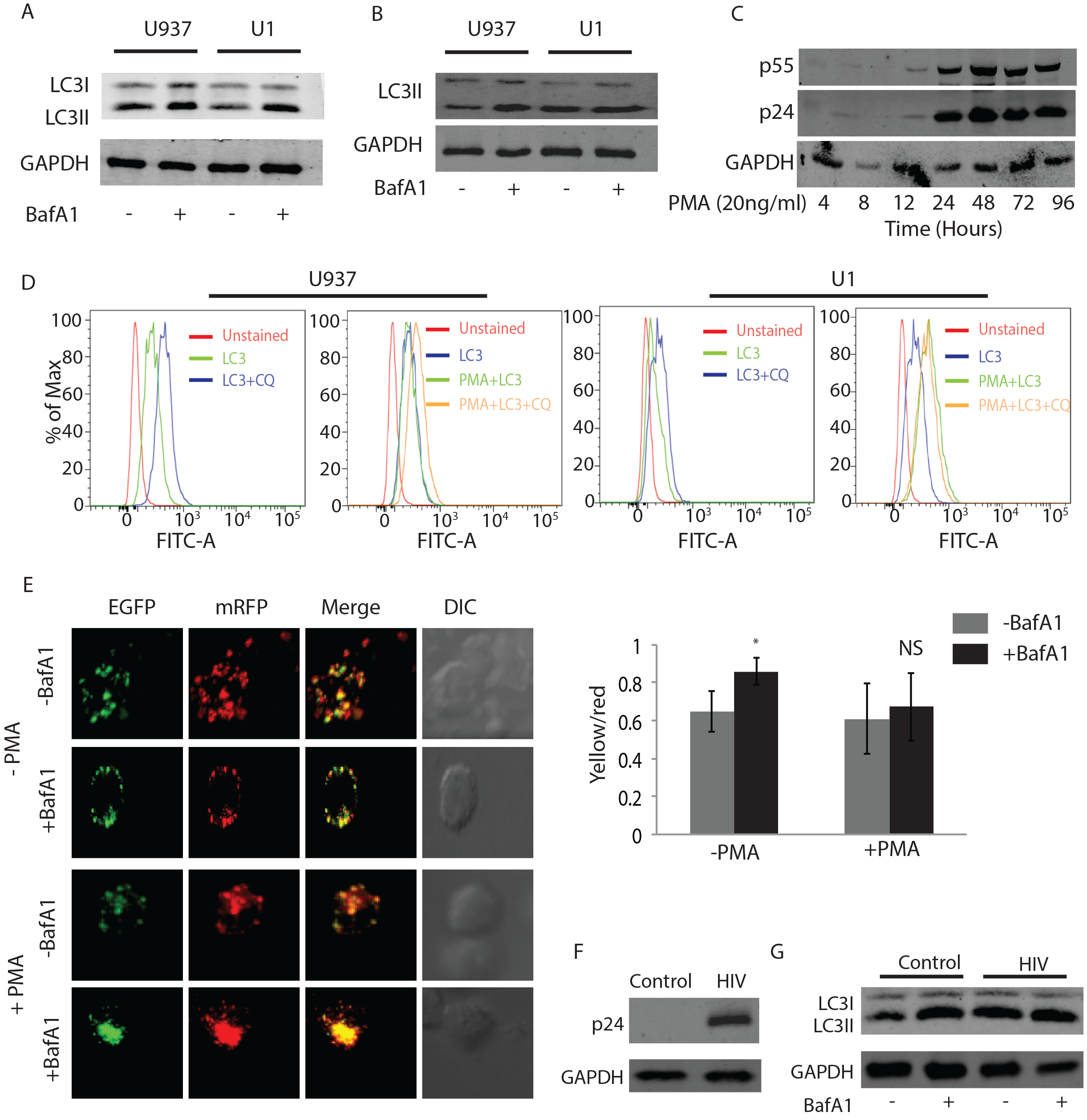
Replication of HIV-1 in macrophages results in autophagy block. Cell lysates of U937 and U1.1 monocytes without PMA treatment (A) or with 20ng/ml PMA treatment for 48 hours (B) were prepared from cells, in the absence or presence of Bafilomycin A1 (BafA1, 100nM) and immune-blotted for LC3B. Baf A1 was added three hours before the time point as described in materials and methods. (C) U1.1 cells were treated with PMA and HIV-1 activation was monitored using p24 immuno-blotting at indicated time points post PMA treatment. (D) U937 and U1.1 monocytes (without PMA treatment) or macrophages (with PMA treatment) were stained for endogenous LC3B upon mild permeabilization using saponin (conc.) and flow cytometry was performed to monitor autophagy flux in the absence or presence of autophagy inhibitor chloroquine (100nM) at 48 hours. (E) U1.1 cells were transfected with RFP-EGFP-LC3 construct and cells were kept with or without PMA (20ng/ml) after transfection for 48 hours. Baf A1 was added 3 hours before the time point. Yellow to red puncta of LC3B were quantified for autophagy flux usingImaris software (see materials and methods). (F) PMA differentiated U937 macrophages were infected with HIV-1 (pNL4-ADA8) at 1:0.01 MOI and p24 levels were probed by immune-blotting in the total cell lysates generated at 48 hours post-infection. (G) In the above experiment, HIV-1 infected U937 cells were either treated with BafA1 or left untreated during final three hours of the assay time (48 hours). LC3 levels were probed in control and HIV-1 infected U937 cells in BafA1 treated or untreated sets by immuno-blotting.

### Productive infection of U937 macrophages with HIV-1 results in impairment of autophagic flux

While results in figure 1 show impairment of autophagic flux upon reactivation of latent HIV-1 infection, we next assessed how host autophagy response was regulated during productive infection with HIV-1. PMA-treated U937 macrophages were infected with HIV-1 (pNL-AD8; CXCR5 tropic) at 0.1 MOI using spinnoculation and at 48 hours post-infection, cells were treated with or without BafA_1_ for three hours before lysing the cells and probed for LC3 by immunoblotting. Productive infection was confirmed by immunoblotting from the cell lysates (Fig. 1F). Similar to our data in the activated U1.1 macrophages, we found autophagic flux was blocked in U937 macrophages upon productive infection with HIV-1 as evident by HIV-1-induced increase in LC3-II levels, which did not further increase upon BafA_1_ treatment (Fig. 1G). Thus, our data point to an impairment of autophagic flux both during reactivation of HIV-1 or during productive HIV-1 infection in macrophages.

### Inhibition of autophagic flux in activated U1.1 macrophages or productive HIV-1 infection in U937 macrophages helps better survival of *Mtb* and NTMs strains

Autophagic flux depicts the crucial step of autophagosome maturation that governs the autophagic cargo degradation in the autolysosomes. Several bacterial pathogens are known to inhibit this late step of autophagic flux in order to survive within the macrophages by preventing xenophagy, which is particularly true in the context of *Mtb*^16,26^. We therefore investigated whether the block in autophagic flux, caused by active HIV-1 infection also facilitated *Mtb* survival within the macrophages. Since, the virulent *Mtb* strain H37Rv is known to inhibit xenophagy flux in the infected macrophages to help its survival^16^, we analyzed some non-tuberculous mycobacterial strains (NTMs) like *M. avium* complex (MAC) and *M. fortuitum* (*Mft*), which are known to have relatively poor survival within human macrophages. PMA-activated U937 (used as control without HIV) or U1.1 (chronically infected with HIV) macrophages were infected with H37Rv, MAC or *Mft* at an MOI of 1:10 and their survival were monitored at 48 hours post-infection. It was evident that activation of HIV-1 replication in U1.1 macrophages resulted in better survival/growth of each of the three mycobacterial strains, with respect to the U937macrophage controls (Fig. 2A, B and C). Similarly, productive infection of U937 macrophages with HIV-1 also led to a much better survival of H37Rv, MAC and *Mft* with respect to the uninfected control conditions (Fig. 2D, E and F). Interestingly, H37Rv and HIV showed some mutualism since in H37Rv infected macrophages p24 levels were significantly higher than those cells that were infected with HIV-1 alone, as evident by p-24 specific RT-PCR and ELSA (Fig. 2G and 2H). Likewise, similar phenomenon was observed when productively infected U937 macrophages were infected with MACs or *Mft* (Fig. 2G, 2H and Fig. S1).

**Figure 2:**
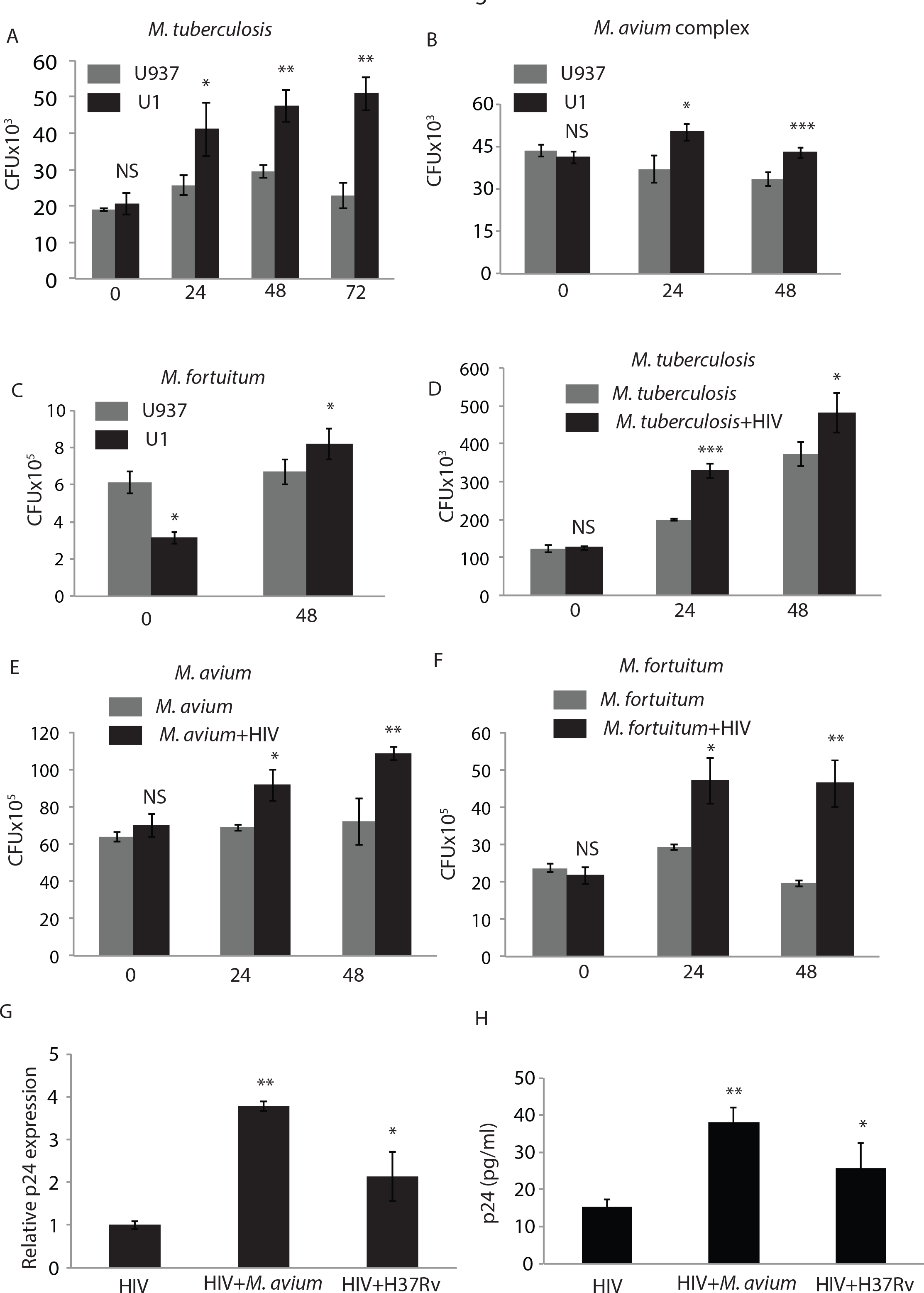
Inhibition of autophagy flux in activated U1.1 macrophages or productive HIV-1 infection in U937 macrophages helps better survival of *Mycobacterium tuberculosis* and non-tuberculous mycobacterial (NTMs) strains. U937 and U1.1 macrophages were infected with either H37Rv (A), *MAC* (B) or *M.fortuitum* (C) at 1:10 MOI and total surviving bacteria were counted at 24 and 48 hours post-infection as CFU/ml. In (D), (E) and (F) U937 cells were infected first with HIV-1 at an MOI of 1:0.01 for 48 hours followed by infection with either H37Rv, *MAC* or *M. fortuitum* respectivelyat MOI of 1:10. Total surviving bacteria were counted at 24 and 48 hours post-infection as CFU/ml. (G) U937 cells were infected with HIV-1 as mentioned above and subsequently infected with H37Rv or *MAC*. Viral replication was measured by monitoring p24 transcript levels in the infected cells at 48 hours post-bacterial infections. (H) In the same assay as above, ELISA was performed to calculate the viral p24 titres in the culture supernatants at 48 hours post bacterial infection (H37Rv and *MAC*), *p-value<0.05; **p-value<0.005; ***p-value<0.001.

### Xenophagy flux is inhibited in HIV-1 infected U937 cells

We previously demonstrated in H37Rv-infected macrophages, that xenophagy (antibacterial autophagy) was selectively impaired without compromising the basal autophagy flux^16^. This enabled better survival of *Mtb* within macrophages^16^. Since, HIV-1 infection too inhibited autophagy flux, we examined whether the increased bacterial survival in HIV-1 infected cells was at least in part due to an additional block in the xenophagy. Xenophagy flux is measured by comparing the localization of bacteria to the LC3^+^ compartment in the absence or presence of BafA_1_ treatment ^25^. As reported earlier, H37Rv-infected cells did not show any xenophagic flux either alone or when co-infected with HIV-1 (Fig. 3A). Additionally, in HIV-1 infected cells, bacterial co-localization to LC3^+^ compartments was much lower compared to non-HIV-1 infected control cells, especially in the presence of BafA_1_ (Fig. 3A). In the context of MACs, there was significant xenophagic flux in the non-HIV-1 condition, which was nearly abolished in the HIV-1 co-infected cells (Fig. 3B). This correlated with the increased survival of MAC in HIV-1 infected U937 macrophages shown in the previous section.

**Figure 3:**
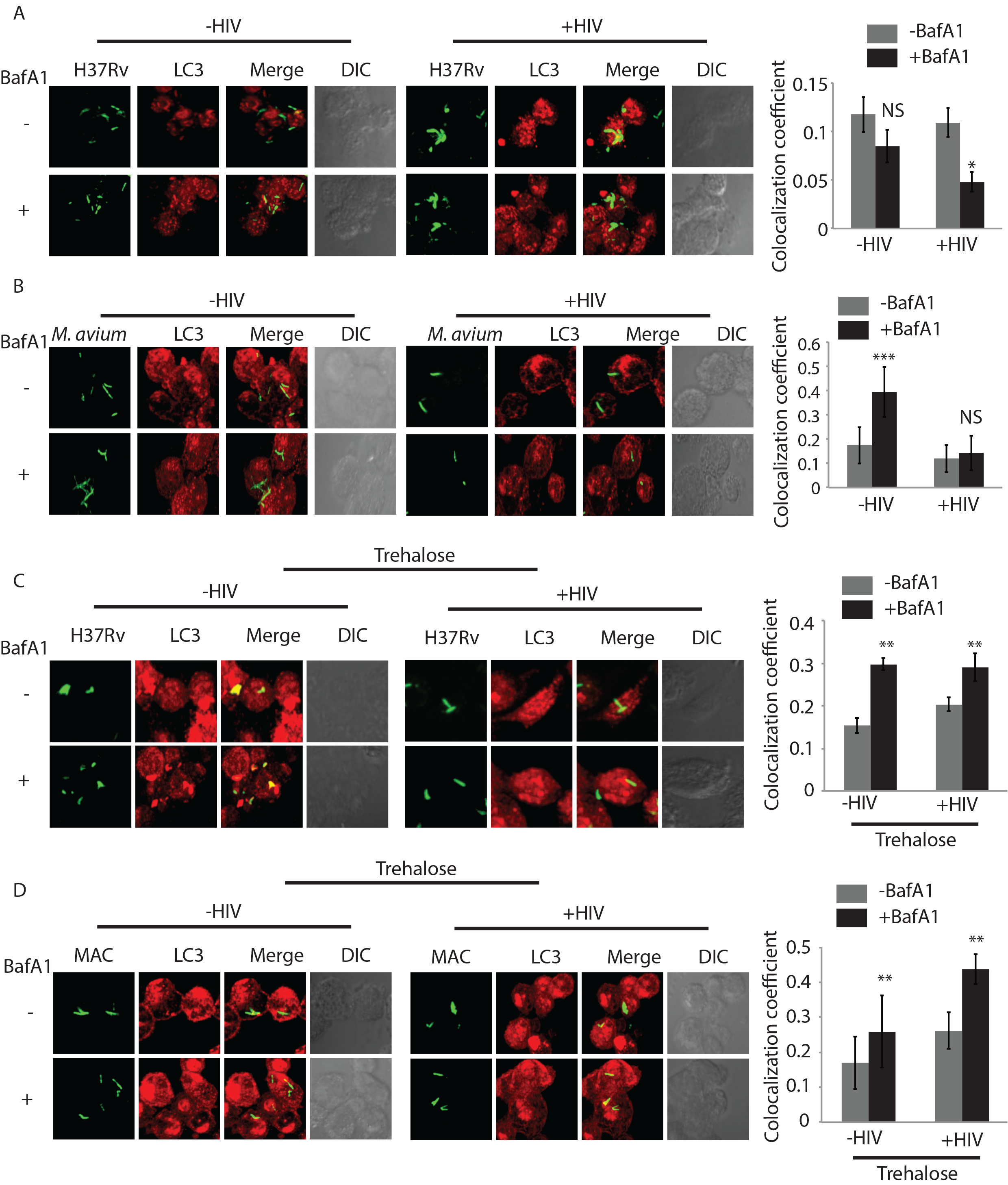
Modulation of xenophagy flux in single and co-infected U937 macrophages in absence and presence of autophagy modulator trehalose. U937 macrophages either uninfected or infected with HIV-1 pNL4-ADA8 (1:0.01 MOI) for 48 hours were subsequently infected with PKH67-labeled (green) H37Rv (A) or *MAC* (B) at an MOI of 1:10. Three hours before the experimental time point (48 hours), one half of the cells were treated with BafA1 (100nM). At 48 hours, cells were fixed and stained with anti-LC3 antibody followed by anti-rabbit-Alexa568 (red) and analyzed under the confocal microscope. Co-localization between bacteria and LC3 compartment in the presence or absence of BafA1 treatment was calculated by analyzing the confocal images in Imaris (see methods, values are mean±SD). For (C) and (D), U937 macrophages either uninfected or infected with HIV-1 pNL4-ADA8 (MOI 1:0.01) for 48 hours were subsequently infected with PKH67 labeled H37Rv or *MAC* (green) respectively at 1:10 MOI. At 36 hours post-bacterial infection, cells were treated or left untreated with trehalose (100mM). Finally before sample harvest at 48 hours, cells from each of the groups were either treated with BafA1 (100nM, 3 hours) or left untreated. At 48 hours, cells were fixed and stained with anti-LC3 antibody followed by anti-rabbit-Alexa568 (red) and analyzed under the confocal microscope. Co-localization between bacteria and LC3 compartment in the presence or absence of BafA1 treatment was calculated by analyzing the confocal images in Imaris (see methods, values are mean±SD; *p-value<0.05; **p-value,<0.005; ***p-value<0.001).

### Xenophagy flux can be induced by the known autophagy inducer trehalose

To further establish that the effect of HIV-1 infection on mycobacterial survival was primarily due to the inhibition of autophagic/xenophagic flux in the host macrophages, we assessed whether stimulation of autophagy could reverse this phenomenon and facilitate bacterial killing. For inducing autophagy, we used known autophagy enhancers, such as, rapamycin, 1, 25-dihydroxy cholecalciferol (Vitamin D_3_) and trehalose. Treatment with either rapamycin or Vitamin D_3_ led to a significant reduction in the bacterial CFU in H37Rv infected macrophages alone, consistent with the existing literature ^15, 17, 27^, (Fig. S2A). Trehalose was included in our study, because it is considered a safer, mTOR-independent autophagy inducer, and has an established role in rescuing the disease phenotypes in mouse models of neurodegenerative diseases ^28,29, 30^ and therefore clinically more acceptable for treatment. Interestingly, trehalose is an important component of the mycobacterial cell wall and it is known to get secreted out of the cell during the synthesis of cell-wall components. Trehalose transporters present on the *Mtb* cell wall take the secreted trehalose back, which is important for *Mtb* virulence and survival ^22^. Consistent with its known pro-autophagy function, trehalose treatment resulted in dramatically reduced bacterial load in H37Rv-infected U937 macrophages (Fig. S2A). During *in vitro* cultures, presence of excess trehalose did not interfere or increase bacterial growth (Fig. S2B and S2C). We confirmed that trehalose indeed induced high autophagy flux in U937 macrophages (Fig. S2D), which was even higher than starvation-induced autophagy flux in these macrophages (Fig. S2D). We next measured xenophagy flux in H37Rv-infected cells or in H37Rv-HIV-1 co-infected cells upon trehalose treatment. Trehalose treatment caused substantial H37Rv-specific xenophagy flux, as evident from increased bacterial co-localization to LC3^+^ compartment in the presence of BafA_1_ (Fig. 3C). Trehalose-induced H37Rv-specific xenophagy flux was equally significant in the absence or presence of HIV-1 coinfection (Fig. 3C). Similar effects of trehalose were seen with MAC (Fig. 3D). Thus, trehalose facilitated xenophagic flux of infecting mycobacteria in the host macrophages.

### Trehalose-induced Xenophagy flux enables the killing of mycobacterial species

Since trehalose treatment could induce both autophagic and xenophagic flux in macrophages, irrespective of the presence or absence of HIV-1 co-infection, we next assessed intracellular survival of *Mtb*, MAC and *M. fortuitum* under these conditions. Trehalose treatment led to a significant decline in the intracellular survival of H37Rv, MAC and *Mft* in U937 macrophages as evident by the reduced intracellular bacterial load (Fig. 4A, B and C). Likewise, trehalose exerted similar effects in enhancing bacterial killing in HIV-1 infected U937 macrophages, where bacterial survival is otherwise more favorable (Fig. 4A, B and C). The effect of trehalose treatment on bacterial survival was robust since both pre-activation or treatment with trehalose post-infection was equally effective in reducing the bacterial survival within U937 macrophages (Fig. S2E).

**Figure 4:**
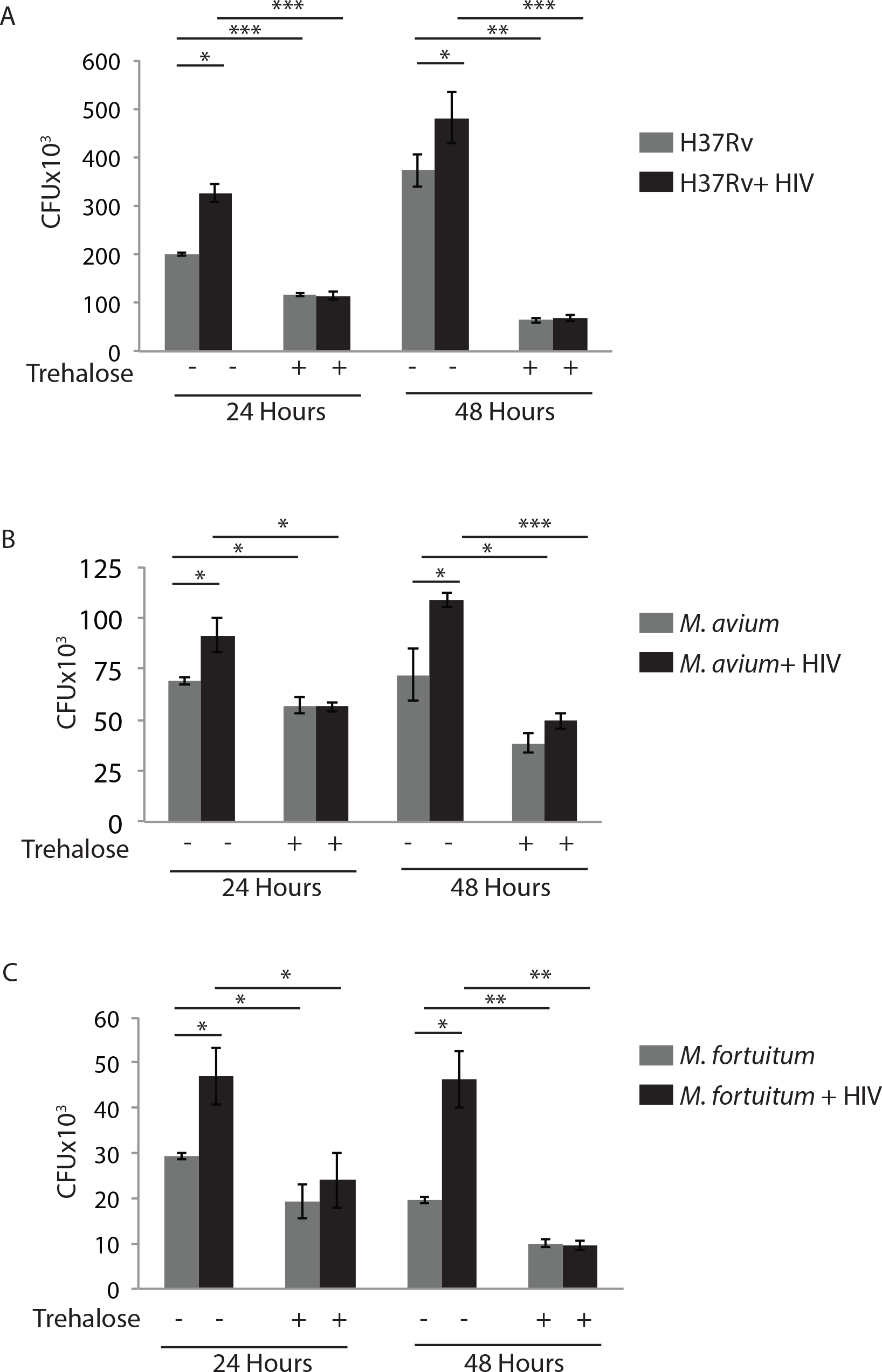
Trehalose induced Xenophagy flux helps in killing kill mycobacterial species. U937 macrophages uninfected or infected with HIV-1 pNL4-ADA8 (MOI 1:0.01) for 48 hours were subsequently infected with H37Rv (A), *MAC* (B) or *M.fortuitum* (C) at the MOI of 1:10. For each group of infections, samples were subsequently either treated with trehalose (100mM) or left untreated for the final 12 hours and cells were harvested at 24 or 48 hours post bacterial infection. Cells were lysed and plated for CFU counting. Values represent mean±SD, from nearly three independent experiments; *p-value<0.05; **p-value<0.005; ***p-value<0.001.

### Trehalose treatment results in lysosomal Ca^2+^ release and activation of TFEB

Trehalose is known to induce autophagy in an mToR-independent manner ^18^. Recently trehalose was shown to inhibit glucose transporters at the cell surface, specifically GLUT3 and GLUT8, thereby causing a pseudo-starvation like response to induce autophagy^19, 20^. However, more recently it was also reported that trehalose can enter the host cell through “GLUT8” transporter and may have intracellular targets for autophagy induction ^21^. Since trehalose treatment in our experiments was for 12 hours or more, we hypothesized that an early increase in autophagy may be followed by increased synthesis of autophagy-and lysosome-related proteins. The increased expression of genes related to autophagy and lysosome function is dependent on the activation of TFEB, a key transcription factor upstream to the autophagy pathway and lysosomal biogenesis genes^31–33^. Using immuno-staining, we confirmed trehalose treatment in U937 macrophages led to a substantial increase in the translocation of TFEB to the nucleus (Fig 5A). Activation of TFEB requires its dephosphorylation by calcineurin, a serine-threonine phosphatase that is activated by intracellular Ca^2+^ ^34^. The release of Ca^2+^ from lysosomal lumen is an important signal for activation of calcineurin ^34^. While the elevation in intracellular Ca^2+^ under any condition is easy to monitor using intracellular Ca^2+^ dyes, we wanted to ascertain whether trehalose causes lysosomal Ca^2+^ release. To that end we used the Ca^2+^-dependent FRET reporter Yellow-chameleon which is fused with the LAMP1 lysosomal localization signal (LAMP1-YCAM)^35^. We transfected HEK 293T cells with LAMP1-YCAM and at 36 hours post-transfection, treated these cells with trehalose for another 12 hours. In the instance of increased Ca^2+^ release from lysosome the LAMP1-YCAM traps the Ca^2+^ and show increased FRET signals. In trehalose-treated cells, we observed enhanced FRET signal from the LAMP1-YCAM reporter suggesting increased Ca^2+^ release from the lysosome (Fig. 5B). The cytosolic YCAM also showed increased FRET signal in trehalose-treated cells (Fig. S3A). The lysosome localized channel protein TRPML1 or mucolipin regulates the release of Ca^2+^ from the lysosome ^34, 36^. We therefore performed siRNA-mediated knock down of TRPML1 and monitored the LAMP1-YCAM FRET signals in trehalose-treated cells (Fig 5E). More than 60% knockdown of TRPML1 was achieved in U937 cells (Fig S3B). In TRPML1 knockdown cells, there was no increase in LAMP1-YCAM FRET signal upon treatment with trehalose (Fig. 5C). Consistent with this data, trehalose dependent nuclear localization of TFEB was completely abolished in TRPML1 knockdown cells (Fig. 5D). The loss in TFEB nuclear translocation also led to a decline in trehalose-mediated autophagy induction in these cells (Fig. 5E). Together our data reveal a new mechanism of autophagy induction by trehalose in macrophages via lysosomal Ca^2+^ release and TFEB activation.

**Figure 5:**
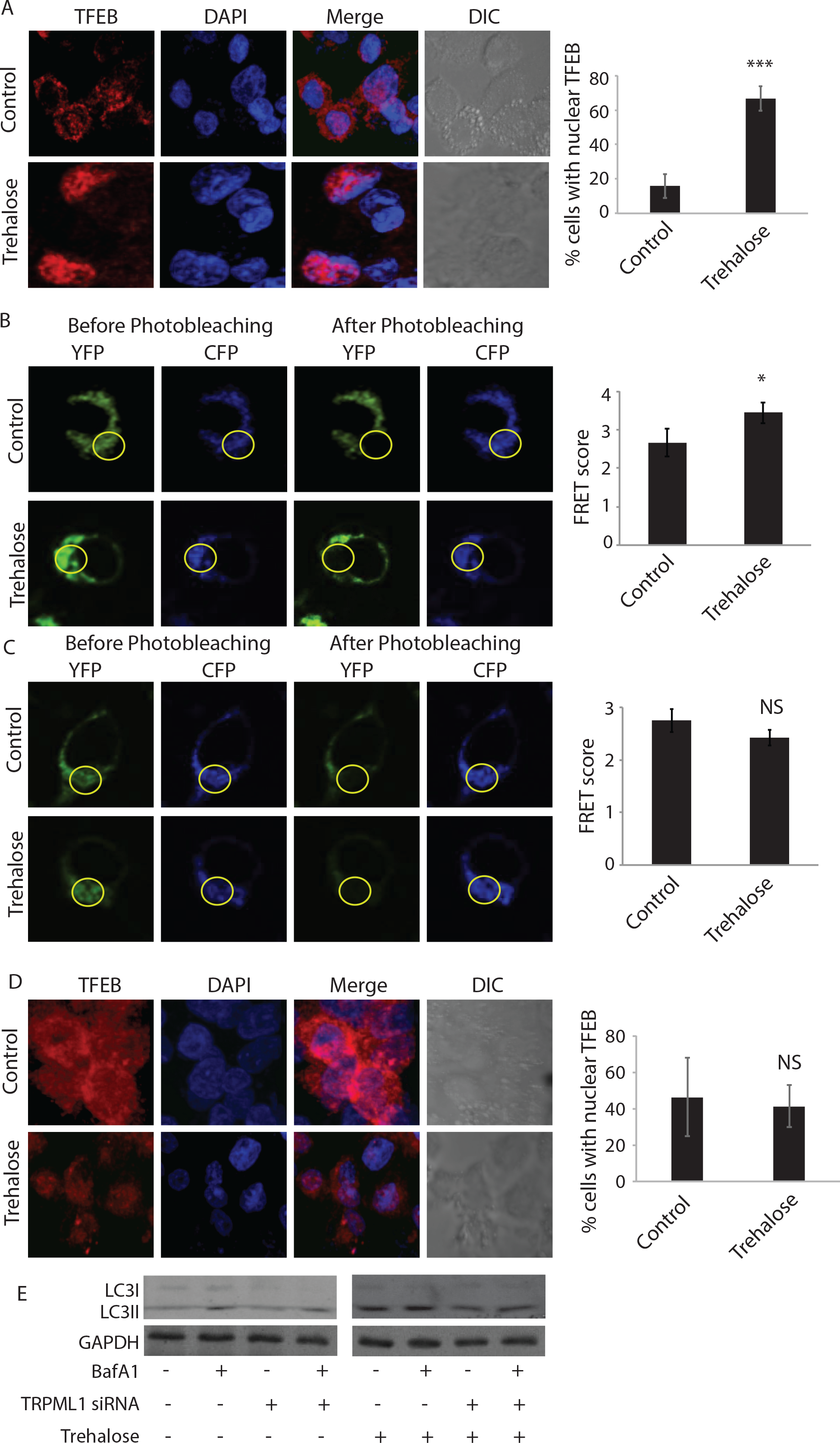
Trehalose treatment results in Lysosomal Ca^2+^ release dependent activation of TFEB. (A) PMA treated (20ng/ml) U937 macrophages either untreated or treated with trehalose (100mM, 12 hours) were fixed and stained with anti-TFEB antibody followed by anti-rabbit Alexa 568 (Red) and DAPI (Blue) and images were acquired under confocal microscope. Percent cells showing nuclear translocation of TFEB was calculated by manually counting the cells in Imaris software. (B) HEK293T cells were transfected with LAMP1-YCAM. At 36 hourspost-transfection trehalose (100mM) was added for 12 hours, cells were fixed and images were acquired under confocal microscope using FRET module. Ratio of CFP to YFP fluorescence before and after photo bleaching is depicted as FRET scores at the right (see methods). For (C) HEK293T cells transfected with LAMP1-YCAM were treated with TRPML-1 siRNA (200nM for 36 hours) or control siRNA. Subsequently, cells were treated with trehalose (100mM, 12 hours) and FRET score was calculated under TRPML1 knock down condition as described above. (D) PMA treated (20ng/ml) U937 macrophages were first treated with TRPML1 or control siRNA for 36 hours, followed by treatment with trehalose (100mM, 12 hours). Cells were fixed and stained with anti-TFEB antibody followed by anti-rabbit Alexa 568 (Red) and DAPI (Blue) and images were acquired under confocal microscope. Percent cells showing nuclear translocation of TFEB was calculated by manually counting the cells in Imaris software. (E) PMA treated U937 macrophages were treated withTRPML-1 or control siRNA. At 36 hours of siRNA transfection, cells were treated with trehalose (100mM, 12 hours). Cell lysates were generated keeping a BafA1 treated set and untreated set for each of the siRNA groups and were probed for LC3 using immunoblotting; *p-value<0.05; **p-value<0.005; ***p-value<0.001.

### Trehalose serves as phosphatidylinositol (3,5) bis-phosphate [PI(3,5)P_2_] agonist to activate TRPML1

How trehalose triggers lysosomal Ca^2+^ release via TRPML1, as shown above, is not clear. Activation of TRPML1 depends on PIKfyve, a class I PI3Kinase, which enzymatically converts PI3P into PI (3,5)P_2_^37^. PI(3,5)P_2_ thus generated serves as TRPML1 agonist and activates the channel for Ca^2+^ release from the lysosomal lumen ^38^. We found that siRNA-mediated knockdown of PIKfyve resulted in a loss in trehalose-induced lysosomal Ca^2+^ release and TFEB localization to the nucleus (Fig. S4). To further ascertain a role of PI(3,5)P_2_ in trehalose-induced autophagy, we transiently expressed an EGFP construct fused to PI (3,5)P_2_ interacting domain of TRPML1 in HEK293T cells^39^. We found a significant increase in PI(3,5)P_2_ fluorescence/punctas in cells treated with trehalose compared to the untreated control (Fig. 6A, B) suggesting that trehalose acts as a PI(3,5)P2 agonist to activate TRPML1.

**Figure 6:**
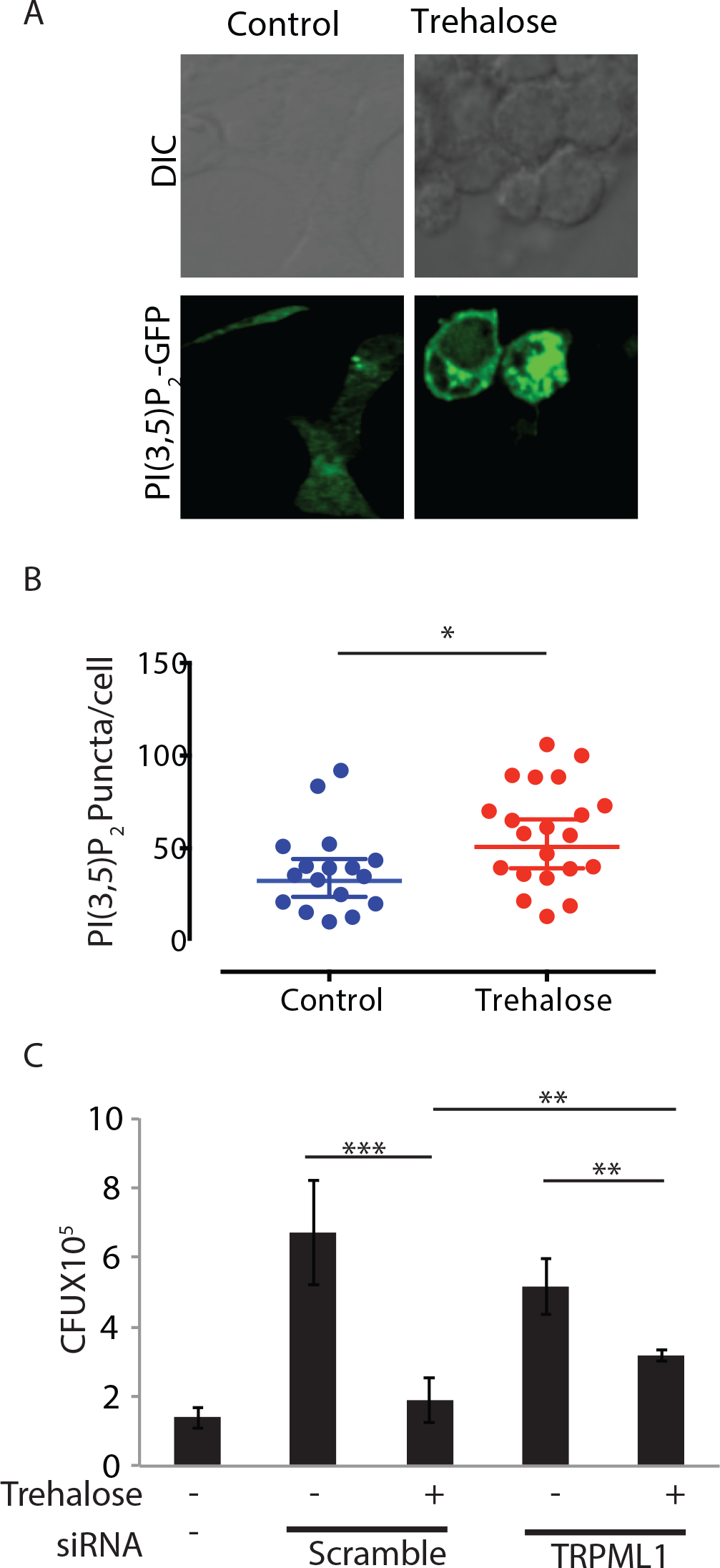
Trehalose serves as Phosphatidyl Inositol (3,5) bis-phosphate [PI(3,5)P_2_] agonist to activate TRPML1. (A) HEK293T cells were transfected with mEGFP-[PI(3,5)P_2_] and after 36 hours were treated with trehalose (100mM) for additional 12 hours. Cells were fixed and images were acquired using confocal microscopy. (B) [PI(3,5)P_2_] puncta/cell were counted in untreated and trehalose treated samples by Imaris software and presented as the dot plots (see methods, *p<0.05). (C) PMA differentiated U937 macrophages were infected with H37Rv at 1:10 MOI followed by siRNA treatment against TRPML-1 at 6 hours post-addition of bacteria. Cells were harvested at 48 hours post infection, lysed and plated to get total CFU count. Values represent mean±SD from two independent experiments; *p-value<0.05; **p-value<0.005; ***p-value<0.001.

### TRPML1 knockdown rescues trehalose-mediated killing of *Mtb* in U937 macrophages

Having established the lysosomal Ca^2+^ release via TRPML1 as the critical pathway for trehalose-mediated autophagy induction, we next wanted to see whether TRPML1 was indeed essential for trehalose-mediated killing of mycobacteria in U937 macrophages that we have demonstrated (Fig. 4A-C). We found that while treatment with trehalose led to a significant decline in H37Rv CFU, in U937 macrophages transfected with scrambled siRNA, siRNA mediated TRPML1 knockdown partially but significantly prevented this effect of trehalose on mycobacterial killing (Fig. 6C). This suggests that the mode of autophagy induction by trehalose, identified in this study, is relevant for its effect on bacterial xenophagy.

### Trehalose treatment results in reduced HIV load in PBMCs from HIV patients

Our data show that while co-infection with HIV-1 rescued the bacterial killing in the macrophages, the reverse was also true, since p24 levels were increased in the coinfected macrophages (Fig. 2). We therefore examined whether trehalose treatment could also directly impact on the HIV-1 replication or survival. PBMCs isolated from healthy volunteers, were infected with clinical strains of HIV-1 subtype C at 200x and 1000x of TCID_50_. HIV-1 infected PBMCs were cultured in the presence or absence of trehalose for 7 days and p-24 in the supernatant was measured on day 3 and day 7 by ELISA. At both the doses of HIV-1 and the time-points studied, treatment with trehalose led to massive decline in the p-24 levels in the supernatant (Figs. 7A and 7B). In PBMCs, where lymphocytes outnumber the monocytes, we analyzed whether trehalose was also effective on the lymphocytes. In the samples from which we performed p24 ELISA using supernatant, we stained the cells with p24 and CD4 and analyzed by FACS. It was evident that in the CD4^+^ cells, treatment with trehalose resulted in a decline in p24 staining (Fig. S5A and S5B). Having observed the effect of trehalose on *ex vivo* infection model, we finally investigated whether trehalose could also impact survival/replication of HIV-1 in PBMCs of HIV-1 infected donors. We recruited 4 treatment-naïve HIV-1 infected donors and cultured their PBMCs for 7 days in the presence or absence of trehalose. Strikingly trehalose treatment was also able to inhibit HIV-1 replication/survival in these clinically relevant cellular platforms (Fig 7C). Taken together, our data implies that trehalose is an effective autophagy inducer for the clearance of HIV-1 and mycobacterial species.

**Figure 7:**
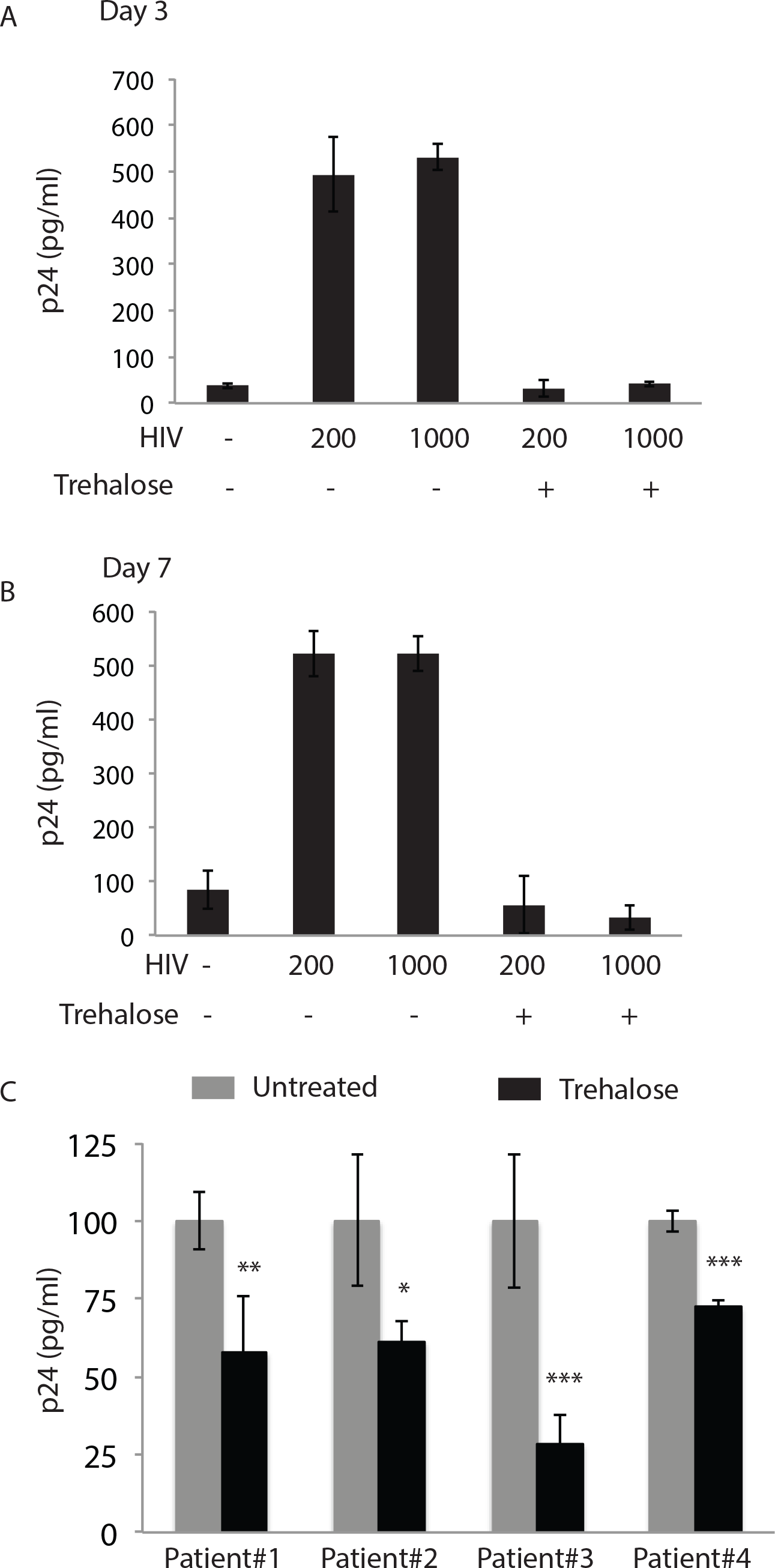
Trehalose treatment results in reduced viral load in PBMCs from HIV-1 infected donors. PBMC from seronegative healthy donors were isolated and infected with clinical isolate of HIV-1 subtype C at doses of TCID_200_ and TCID_1000_ for 24 hrs. Trehalose (100mM) was added post infection and supernatants were collected on Day 3 (A) and Day 7 (A) post treatment for p24 ELISA. Values are mean±SD from duplicates. (C) Infected PBMCs isolated from four ART-naive HIV-infected donors were treated with trehalose and supernatant ELISA for p24 was performed at day 7. Values are mean±SD from triplicates; *p-value<0.05; **p-value<0.005; ***p-value<0.001.

## Discussion

This study was designed to address whether the host innate defense mechanisms perturbed upon HIV-1 infection inadvertently ends up facilitating opportunistic bacterial pathogens like *Mtb* and other non-tuberculous mycobacterial strains. The innate defense pathway of interest in this study is autophagy, which has been extensively characterized during pathogenic bacterial infections including *Mtb* infections^17, 27, 40^. Autophagy regulation has also been explored in the context of several viral infections including HIV-1, dengue and influenza among others^41^. In case of HIV-1, it was initially reported that while its infection results in a surge in the number of autophagosomes formed, it however impairs the maturation of autophagosomes and blocks its fusion with the lysosomes, which was subsequently reported to be dependent on the viral accessory protein Nef ^42, 43^. Therefore our finding that either upon activation of latently integrated HIV-1 in U1.1 cells or by infecting U937 macrophages with HIV-1 caused a block in autophagy flux is consistent with the literature. In different cells including activated macrophages, a block in autophagy flux could result in the accumulation of damaged and depolarized mitochondria, leading to increased cellular ROS and induction of apoptosis^44^. However upon HIV-1 infection, while autophagic flux was suppressed, there was no mitochondrial depolarization observed in these infected cells (Fig. S6). Curiously, it is known that HIV-1 Tat, another accessory protein of HIV-1, can localize to the mitochondria and ensure its hyperpolarized state by modulating the Ca^2+^ channel activities^45^. Interestingly, recent studies show that slight increase in cellular glutathione redox state may activate latent HIV-1 infections into active replicative forms ^23^. As our data show, autophagy flux gets inhibited upon activation of HIV-1 replication in U1.1 cells, ensuring mitochondria in hyperpolarized state would certainly help prolong host cell survival. The role of autophagy in targeting intracellular pathogens to the lysosomes for degradation is well-established^9, 16, 26^. Our results, for the first time report that in cells with active HIV-1 replication, *Mtb*, MAC and *Mft* survived better than the control cells. How bacterial infection in macrophages helped increased p24 levels during co-infection experiments, however, is not very apparent. Previously it was reported that co-infection with *Mtb* results in increased HIV-1 viraemia^3^. Increased HIV-1 replication in co-infected cells was possibly independent of any direct effect on autophagy since we know that H37Rv selectively inhibits only xenophagic flux without impacting the basal autophagic flux whereas for NTMs like MACs or *Mft* there are no such reports suggesting block in autophagic flux or xenophagic flux. At least our experiments show that in MAC-infected macrophages there is no block in xenophagic or autophagic flux.

Using autophagy modulators to target both HIV-1 and *Mtb* has been explored earlier, where it was shown that vitamin D_3_ could kill both *Mtb* and HIV either alone or under co-infection conditions ^13–15^. However, so far, there is no study directly establishing HIV-1 mediated block in autophagy as the causative mechanism behind opportunistic bacterial infections. Using non-tuberculous mycobacterial strains in this study, we are able to demonstrate that the usually non-pathogenic strains also can benefit from the HIV-1 induced autophagy block and therefore could cause opportunistic infections. In fact virulent *Mtb* itself can inhibit xenophagy flux, without perturbing the homeostatic autophagy flux in the cells^16, 26^. This aspect is evident in our results on xenophagy flux, where H37Rv localization to LC3^+^ compartment did not increase in the presence of BafA1 in the absence or presence of HIV-1 infection. In the presence of HIV-1, H37Rv localization to LC3^+^ compartment rather declined in the BafA_1_ treated set. It is not apparent why in HIV-1 infected cells, *Mtb* co-localization to LC3^+^ compartment would decline in the presence of BafA_1_. It nonetheless raises a question on whether the increased bacterial survival observed in our experiments, at least in the case of H37Rv infection was indeed due to impairment of autophagy flux by HIV-1. It is plausible that any residual xenophagy flux occurring during H37Rv infection gets more strongly or entirely blocked in the presence of HIV-1, which possibly is beyond the detection limit of our experimentation. Alternatively, in the context of HIV-1 induced increase in H37Rv survival, improved mitochondrial health possibly plays a more defining role than HIV-1 induced autophagy block. Nevertheless, the fact that mTOR dependent or mTOR independent autophagy inducers like vitamin D3, rapamycin and trehalose could kill H37Rv as efficiently as MAC and *Mft* in HIV-1 co-infection conditions, affirms that autophagy indeed is a major player in intracellular bacterial survival even during co-infection with HIV-1.

We focused on trehalose for inducing autophagy because of multiple reasons: a) its ability to induce autophagy is being explored currently for future clinical trials against neurodegenerative diseases associated with accumulation of cytosolic protein aggregates due to dysfunctional autophagy^29,30^; b) it is a natural disaccharide and nontoxic; c) it was more efficient in reducing bacterial load compared to standard inducers like rapamycin and vitamin D_3_. Interestingly, trehalose is an important metabolite for *Mtb*, which is secreted out of the cell during mycolic acid biosynthesis by antigen 85B as the byproduct^22^. The secreted trehalose must be taken back by the bacteria to ensure continued synthesis of cell-wall components. The uptake of extracellular trehalose is regulated by a sugar transport operon of *Mtb* consisting of genes like lpqY, sugA, sugB and sugC^22^. The same study also suggested that inability to recycle trehalose due to loss of sugA operon genes resulted in compromised virulence and intracellular survival however there was no effect on *in vitro* growth of the mutants^22^. It is possible that due to loss of trehalose transporters from *Mtb*, trehalose thus excluded out in the infected macrophages, may induce autophagy in the host cells, thereby killing the bacterium more efficiently. Experimentally proving this hypothesis, however, will be technically challenging for reasons like quantifying the amount of trehalose secreted by mutant strains lacking trehalose transporter. Moreover, there is no apparent intracellular target known for trehalose, making it even difficult to establish the role of trehalose released from intracellular *Mtb* for inducing autophagy. Trehalose was initially reported as an inducer of autophagy through a mechanism that was independent of mTOR regulation^18^. Although the mechanism of action was unclear, it has been recently reported that trehalose causes pseudo-starvation like phenotype by competitively inhibiting GLUT transporters at the cell surface, resulting in activation of autophagy^19^. Starvation induced autophagy however is known to rely on inhibition of mTOR signaling, in addition to activation of AMPK pathway ^46, 47^. Surprisingly, the same group subsequently reported that at least in certain cell types, GLUT8 may serve as trehalose transporter and trehalose may actually have some intracellular targets for inducing autophagy^21^. Given our new findings in this study, it is possible that trehalose could influence autophagy through multiple cellular pathways.

Since the duration of treatment with trehalose in our study ranged between 4 to 48 hours, we opined that trehalose must also increase the autophagy response through transcriptional activation of autophagy-and lysosome-related genes. This assumption led us to study the nuclear translocation of TFEB and the activation of TRPML1 channel on the lysosomes. Incidentally, cytosolic sequestration of TFEB by both HIV-1 and *Mtb* has been reported earlier^48, 49^, therefore trehalose-mediated nuclear translocation of TFEB seemed to just reverse the modulation of this key pathway in autophagy regulation. Trehalose-mediated nuclear translocation of TFEB and autophagy activation in macrophages was also recently implicated as potential therapy for atherosclerosis^50^. Activation of TRPML1 channel is dependent on its interaction with PI(3,5)P_2_, a product of the enzymatic action of PIKfyve ^38, 51^. Role of PIKfyve in regulating autophagy in itself is debatable, with literature references supporting both pro-as well as anti-autophagy properties. Thus, while PIKfyve through PI(3,5)P_2_ can activate TRPML1^38, 51^, it is also reported that both PI(3,5)P2 as well as PI3P can activate mTOR pathway, which should negatively regulate autophagy ^52, 53^. Studies also show that PI(3,5)P2 serves as the precursor for entire cellular PI3P content, most likely through some phosphatase activity^37^. However, there is no contradiction on the requirement of PI(3,5)P_2_ in the activation of TRPML1 channel. We therefore used PI(3,5)P_2_ binding domain ofTRPML1 fused with EGFP to conclusively show that trehalose acts as an PI(3,5)P_2_ agonist, which explains the autophagy-inducing property of this molecule. Since TRPML1 knockdown resulted in reduced autophagy flux upon treatment with trehalose, and rescued H37Rv from trehalose-induced killing within the macrophages, our data further strengthens the direct role played by autophagy in the bacterial killing, either alone or in co-infection with HIV-1.

The most fascinating and clinically relevant finding in this study however was the effect of trehalose treatment on HIV-1 replication during *ex vivo* infections as well as in the PBMC cultures derived from treatment naïve HIV-1 infected donors. Substantial decline in p24 that we have found in the above conditions suggest the viral killing ability of trehalose-induced autophagy. Since in PBMC cultures, majority of cells are lymphocytes, it is evident that trehalose-induced autophagy can not only clear HIV-1 infection in monocyte/macrophages but also in CD4^+^ T cells. The ability of trehalose to induce autophagy and control bacterial and viral infections in different immune cells makes it a promising candidate for future management of HIV-1 and associated opportunistic infections. Furthermore, HIV-1 induced neuropathy is known to get manifested through inappropriate autophagy response^54^. Since trehalose has been shown to be beneficial in the transgenic models of neurodegenerative disorders, there is an immense potential of this disaccharide in the management of HIV-1 infection and disease progression, which must be further explored in combination with ART.

In conclusion, we demonstrate that HIV-1 mediated inhibition of autophagy is an important regulator of opportunistic bacterial infections. Furthermore we show that trehalose, a naturally occurring disaccharide, can induce autophagy and kill *Mtb* and NTMs alone or during co-infection with HIV-1. Interestingly, we also delineate an entirely novel mechanistic aspect of trehalose-mediated autophagy induction and show that it serves as a PI(3,5)P_2_ agonist. Finally, the ability of trehalose to kill HIV-1 in PBMCs from healthy volunteers infected *in vitro* or from treatment naïve HIV-1 infected donors show tremendous potential of this molecule in combating HIV-1 and associated pathogenesis.

## Author Contributions

Performed experiments: VS, MM, PK; Analyzed data: VS, DK, Provided reagents: NK, SS (AIIMS), SS; Experiment designs: VS, KL, SS, DK, Wrote manuscript: VS, SS, DK; Overall conceptualization, supervision and funding: DK.

## Acknowledgements

This study was supported by “Creative and Novel Ideas in HIV Research” (CNIHR) award from National Institutes of Health (NIH)-Centers for AIDS Research (CFAR) in association with International AIDS Society (IAS) (Grant No. 5P30AI027767–27 to DK). VS is a recipient of Senior Research Fellowship from Council of Scientific and Industrial Research, Govt. of India. SS is funded by Wellcome Trust Seed Award (109626/Z/15/Z) and Birmingham Fellowship. DK and SS are also jointly funded by UKIERI-DST Thematic Partnership Award (2016-17-0087). We thank Murali K Kaja for his support to CNIHR grant and N. Siddappa for providing U1.1 cells. We acknowledge Shahid Jameel for providing infectious clones and cell lines. We acknowledge support from Department of Biotechnology, Govt. of India for supporting TACF (Tuberculosis Aerosol Challenge Facility), the BSL3 facility.

## Materials and Methods

**Ethics Statement**: All the experiments with PBMCs, either from healthy volunteers or HIV-infected donors, were conducted at AIIMS, New Delhi. The protocol was approved by the institutional ethics committee of AIIMS, New Delhi (IEC/NP-295/2011).

### Cell culture, Media and reagents

U937 (Sigma Aldrich), U1.1 (NIH reagents program), HEK293T (Clonetech), TZM-bl (NIH reagent program). All cell lines were maintained in either RMPI or DMEM (Gibco Laboratories, Gaithersburg, MD, USA) with Fetal calf serum (Gibco) according to the manufactures protocol. All media reagents HEPES, sodium bicarbonate, glutamine etc. were procured from sigma. Transfection reagents (Jet Prime and Jet Prime buffer) were obtained from Polyplus^Tm^, retro-concentrin were obtained from Clonetech Laboratories Inc.

### Inhibitors, Antibodies, Plasmids, Constructs and other reagents

Bafilomycin A1, Rapamycin, 3 Methyl-Adenine, were obtained from Sigma-Aldrich Co (St Louis, MO, USA). Primary antibodies MAP1LC3B, (Cell Signaling Technology and Novus Biologicals), GAPDH (Santa Cruz Biotechnology), TFEB (Bethyl Laboratories, Inc.), Anti HIV-1 p24 (Abcam), anti HIV-1 p24 FITC (Beckman Coulter, Inc.), p70S6 Kinase and phospho p70S6 kinase (Cell Signaling Technology). Anti TRPML-1 (Novus Biologicals). All IR conjugated secondary antibodies were obtained from LI-COR Biosciences (Lincoln, NE, USA) and Alexa Flour dye conjugated secondary antibodies were procured from Invitrogen Molecular Probes, Carlsbad, CA, USA). Phorbol 12-myristate 13-acetate, (PMA), PKH67 dye, β-galactosidase, DMSO, polybrene (1,5-dimethyl-1,5-diazaundecamethylene polymethobromide, hexadimethrine bromide), BSA (Bovine Serum Albumin), MTT (1-(4,5-Dimethylthiazol-2-yl)-3,5-diphenylformazan) and paraformaldehyde, were AMRESCO Inc. JC-1 (Thermo Fischer SCIENTIFIC), ptfLC3 was a kind gift from Sovan Sarkar (Addgene-Plasmid # 21074), YC (YCaM) 3.6 was obtained from Addgene (Addgene-Plasmid # 58182), and LAMP-1 YCaM was a kind gift from Peter Haynes. mEGFP-PI (3,5) P_2_ (Addgene-Plasmid # 92419) were a kind gift from Geert van den Bogaart.

### Mycobacterial cultures and HIV-1 clones

HIV-1 clones (pNL-AD8) were obtained from Dr. Shahid Jameel, mycobacterial strain H37Rv was obtained from the Colorado State University, Ft. Collins, Co (USA). Clinical strains of *Mycobacterium aviuum* Complex and *Mycobacterium fortuitum* were obtained from All India Institute of Medical Sciences, New Delhi. All mycobacterial cultures were grown in middlebrook 7H9 broth (HiMedia laboratories). HIV-1 infected PBMCs from clinical samples for PBMC experiments were obtained from Department of Biochemistry, AIIMS.

### Cell culture

Human promonocytic cell lines U937 and U1.1 were maintained in RPMI 1640 with 10% FCS at 37C in 5% CO2, humidified incubator. Similarly, HEK293T and TZM-bl cells were maintained in DMEM and 10% FCS. For seeding of U937 and U1.1, cells were treated with 10ng/ml of PMA for overnight and washed subsequently for removing PMA the next day and kept for differentiation for 24 hours. These cells were proceeded for setting up infection experiments. For seeding of HEK293T and TZM-bl, cells were seeded in tissue culture treated Petri-plates or six well plates in complete DMEM media overnight and transfection and titration experiments were carried out respectively.

### Transfection experiments and virus preparation

For preparation of HIV-1 viruses, molecular clones of HIV-1 (pNL-AD8) were transfected in HEK293T cells by using Jet prime method. In brief, 10ug of plasmid DNA was taken and diluted with jet prime buffer according to the manufactures protocol. Jet prime reagent was mixed with DNA containing buffer, vortex briefly and incubated for 10 min. After incubation the mixture was added onto the cells and kept for 4-8 hours. After that fresh media was added to the culture and kept for 36-48 hours Viruses were harvested after 48 hours of transfection by collecting the supernatant, which was subsequently filtered with .45micron filter and kept for further use.

### Concentration of viruses

The harvested supernatant was concentrated by mixing it with retroconcentrin and incubated at 4^O^C for overnight. Next day, the supernatant was pelleted at 1500g for 30 min at 4^O^C the pellet was resuspended in filtered 1X PBS and aliquots of viruses were stored in -80 ° Celsius for further use.

### Titration of viruses

For titration experiments, TZM-bl cells were seeded in six well plate at a density of .3 million cells per well. Cells were treated with polybrene (8ug/ml), kept for 10 min, viruses were added onto the cells, at different dilutions, after removing polybrene. Next cells were incubated for 4 hours for virus adsorption and internalization. After incubation cells were washed twice with 1X PBS and fresh complete DMEM was added and again kept in incubator for 36 hours. β galactosidase assay was performed on these cells for titration of viruses. In brief, media was removed, cells were fixed in .05% gluteraldehyde for 10 min and washed subsequently two times with 1X PBS. Working solution of β galactosidase was added (2/3 ml per well) and kept for 2-16 hours (according to the signal) in incubator. Blue cells were counted in different areas in each well and titre was calculated.

### Bacterial cultures and infection experiments

All bacterial cultures were grown in 7H9 media supplemented with 10% ADC (Becton Dikinson) until log phase. Cultures were then used for infection experiments. CFU plating were done on 7H11 agar supplemented with 10% OADC (Becton Dikinson). Single cell suspension of bacterium for infection experiments was used. For obtaining single cell suspension log phase culture of bacteria were passed through 23*7, 26*5, 30*3 times with above mentioned gauge needles. Quantitation of bacteria was done by taking absorption method where 0.6 OD corresponds to 100 million bacteria per ml at 600nm wavelength. Required number of bacteria was added onto the seeded cells in complete media at MOI of 1:10. Similarly, for confocal experiments bacteria was stained with PKH67 dye according to the manufactures protocol and re-suspended into final media and infection was done. Infection experiments were done for 4 hours and to kill the extracellular bacteria amikacin sulphate (Sigma-Aldrich) was added at a final conc. of 200ug/ml for 2 hours. Then, amikacin sulfate was removed and fresh media was added at kept for scoring the required time points. For CFU experiments culture media was removed and cells were lysed in a buffer (7H9) containing .06% SDS and plating was performed at required time points. Cell viability was monitored by MTT assay. In brief, cells were incubated with working concentration of MTT (1mg/ml) and final reading was taken at 645nm.

### HIV-1 infection experiments

In all HIV-1 infection experiments U937 cells were infected with 1:0.1 MOI of pNL-AD8 strain of HIV-1 by incubating the cells with the virus and spinoculated for 1 hr. at 1200rpm. Then cells were kept for 4 hours in the CO_2_ incubator and washed subsequently to remove any unbound virus and kept for 48 hrs. for further experiments.

### Flow cytometry

LC3BII staining: For flow cytometry experiments cells were seeded in 6 well plate at a density of 1 million per well. At different time points cells were scrapped off and for MAP1LC3BII staining experiment cells were pelleted down at 1000rpm and treated with 0.1 % of saponin for 10 min and subsequently blocked in 3%BSA in 1XPBS and incubated with primary antibody for 1 hour in blocking buffer followed by incubation in Alexa-flour 488 conjugated secondary antibody for 1 hour. After incubation cells were washed with 1 XPBS and re-suspended in 1X PBS and samples were acquired in BD FACS Canto II using FACSDiva acquisition software. The data was analyzed using Flow Jo V.

JC1 staining: U937 cells were treated with 10ng/ml of PMA, washed and infected with pNL-AD8 strain of HIV-1 at an MOI of 1:0.1 and kept for 4 hours Virus was removed and cells were washed twice with 1XPBS to remove any unbound virus and kept for 48 hours. Cells were then treated with JC1 stain (2uM) for 20 min in 37 CO_2_ incubator and analyzed with FACS. Analyses were done by using Flow Jo V.

**siRNA transfections**: HEK-293T cells were either left untreated or transfected with siRNA of either TRPML-1(100nM) or PIKfyve (100nM) for 6 hrs. and kept for 36 hrs. Similarly, U937 cells were seeded with 10ng/ml of PMA, washed and transfected with siRNA of TRPML-1 and kept as described above.

### Nucleofection

U1.1 cells were nucleofected according to the manufactures protocol by using Amaxa 2D-Nucleofector ^Tm^. In brief, the cells were nucleofected by suspending the required amount of DNA (ptfLC3) in the provided suspension buffer and nucleofection reagent was added subsequently according to the kit protocol. The mixture was then taken for nucleofection and cells were kept for 12 hours post nucleofection. The cells were either left untreated or treated with 10ng/ml of PMA for 24 hours. BafA_1_ was added for 3 hours before the time point. At time point cells were fixed and mounted for confocal microscopy.

### Immunofluorescence staining and Confocal microscopy

For staining LC3 U937 cells were fixed and permeabilized with 0.25% TX-100 and subsequently blocked in 3% BSA followed by incubation in primary antibody, from Novus Bilologicals (anti-MAP1LC3BII) at the recommended dilution, and washed two times with PBST and further incubated with secondary antibody tagged with Alexa flour 565nm. Similarly, TFEB staining was done as described above. In brief, U937 cells were fixed and permeabilized with TX-100 and blocked in 3% BSA followed by incubation with anti-TFEB primary antibody, washed and subsequently incubated with secondary antibody tagged with Alexa flour 565nm according to the manufacture’s protocol.

Samples were fixed in PFA and subsequently stained with the required primary and secondary antibody after blocking. Immunofluorescence were done in Nikon Ti-E microscope equipped with 60X/1.4 NA planapochromat DIC objective lens. For FRET Experiments, HEK-293T cells were seeded and were subsequently transfected with YC (YCaM)-3.6 addgene Plasmid # 58182 or LAMP-1 YCaM construct and kept for 36 hours. Addition of trehalose (100mM) was done post transfection for 12 hours.

